# Perplexity as a Metric for Isoform Diversity in the Human Transcriptome

**DOI:** 10.1101/2025.07.02.662769

**Authors:** Megan D. Schertzer, Stella H. Park, Jiayu Su, Fairlie Reese, Gloria M. Sheynkman, David A. Knowles

## Abstract

Characterizing the extensive isoform diversity revealed by long-read RNA-sequencing remains challenging. After removal of technical artifacts, existing pipelines apply arbitrary expression thresholds that filter out bona fide transcript structures, misrepresenting diversity and hindering reproducibility. Instead of discarding isoforms, we propose a fundamentally distinct approach to quantifying isoform diversity using *perplexity*–the effective number of isoforms for a gene, derived from Shannon entropy–wherein every isoform, including low-abundance ones, contributes proportionally to a gene’s diversity. Analyzing 124 ENCODE4 PacBio datasets spanning 55 human cell types, we show that perplexity provides interpretable and reproducible isoform diversity measurements across genes, regulatory levels, and tissues.

## Background

The majority of human genes can produce multiple isoforms (Pan et al. 2008; Wang et al. 2008). Early catalogs were built using cDNA cloning and Sanger sequencing, largely through expressed sequence tags (ESTs) (Gerhard et al. 2004; Harrow et al. 2006). Short-read RNA-sequencing later enabled high-throughput discovery of alternative splicing events across cell types and conditions (Harrow et al. 2012; Frankish et al. 2019; ENCODE Project Consortium 2012). However, short reads typically span only a single exon or splice junction, enabling event-level analysis (Li et al. 2017; Shen et al. 2014), but making it difficult to reconstruct full-length transcripts. As a result, isoform-level quantification has relied on probabilistic read assignment to reference transcripts (Trapnell et al. 2010; Bray et al. 2016; Patro et al. 2017), an error-prone process when isoforms are highly similar, gene structures are complex, or reads originate from unannotated isoforms (Sarantopoulou et al. 2021; Zhang et al. 2017; Steijger et al. 2013). Consequently, low-abundance isoforms in short-read datasets represent low-confidence structures and are routinely filtered out using strict expression cutoffs.

Long-read RNA-sequencing (LRS) overcomes these limitations by capturing full-length RNA molecules, providing direct evidence of transcript structures and enabling detection of combinations of splicing and transcriptional events (Sharon et al. 2013; Garalde et al. 2018; Workman et al. 2019; Al’Khafaji et al. 2023). Thus, LRS is fundamentally reshaping our understanding of the human transcriptome, not only by revealing vast numbers of novel isoforms (Monzó et al. 2025; Pardo-Palacios, Wang, et al. 2024; Reese et al. 2023; Glinos et al. 2022; Leung et al. 2021; Humphrey et al. 2025), but also by challenging assumptions about the interpretation of low-abundance isoforms. LRS shifts the primary uncertainty from whether these isoforms are “real” to determining their biological relevance and function.

Filtering choices during analysis, such as transcript per million (TPM) thresholds, minimum isoform usage cutoffs, and sample recurrence requirements, can significantly impact estimates of isoform diversity. Yet, there is currently no consensus on filtering parameters, and the choice of threshold inevitably misrepresents isoform diversity, overestimating for some genes and underestimating for others. Moreover, while technical artifacts can and should be removed bioinformatically, there is no ground truth to distinguish low-abundance biologically meaningful isoforms from biological noise, meaning that no threshold can be objectively justified over another. As a result, reported estimates of transcriptome complexity can vary substantially (Pardo-Palacios, Wang, et al. 2024), complicating biological interpretation and cross-study comparisons.

Instead of refining thresholds and discarding data, we propose a fundamentally distinct approach that considers all detected isoforms to quantify isoform diversity: *perplexity*, the exponential of Shannon entropy. Perplexity condenses isoform ratio distributions per gene into a single value that reflects the effective number of isoforms. It is a specific *Hill number*, a family of diversity measures initially applied to quantify species diversity in ecological communities (Hill 1973; Jost 2006; Leinster 2020). While Shannon entropy has been applied in transcriptomics, these studies use raw entropy values rather than transforming them to Hill numbers (Martínez and Reyes-Valdés 2008; MacArthur and Lemischka 2013; Gandrillon et al. 2021; Sterne-Weiler et al. 2018; Trapnell et al. 2010; Ritchie et al. 2008; Jones et al. 2024). Hill numbers have been applied in metagenomics and molecular ecology (Ma and Li 2018; Alberdi and Gilbert 2019), but to our knowledge have not been applied to transcriptomics.

Here, we apply perplexity to quantify isoform diversity within 124 ENCODE4 PacBio LRS datasets across 55 human cell types (Reese et al. 2023). We show that perplexity, the effective number of isoforms, provides a principled diversity metric that adapts across genes of varying expression and complexity and is robust across replicates. We extend this framework to distinguish diversity at the transcript and ORF-level, capturing the convergence of multiple transcripts that produce the same protein. To facilitate integration of perplexity metrics into existing pipelines, we provide IsoPlex (https://github.com/fairliereese/isoplex), a Python library for calculating perplexity from LRS data. Together, this study establishes an unbiased and generalizable framework for quantifying isoform diversity across complex genes, cell types, and conditions.

## Results

### Perplexity is a Principled Metric to Quantify Isoform Diversity

LRS captures full-length RNA molecules and provides the most comprehensive view of isoform diversity to date. We analyzed ENCODE4 PacBio LRS data (Reese et al. 2023) using a custom pipeline that collapses reads to isoforms based on shared splice junctions, irrespective of precise read start or end site variation (Supplemental Figure 1A; see Methods). Technical artifacts, such as internal priming and fragmented reads, can be confidently identified, and should be removed prior to diversity analysis (see Methods). After artifact removal, the remaining transcript structures represent high confidence isoforms present in the sample, regardless of abundance. At this stage, we propose a fundamentally distinct approach: quantifying isoform diversity across the complete isoform repertoire, rather than removing low-abundance isoforms as assumed biological noise. We apply the Hill number framework, originally applied to species diversity in ecology (Hill 1973; Jost 2006; Leinster 2020). Leinster (2020) proved that Hill numbers are the only diversity metrics that satisfy reasonable theoretical desiderata (Supplemental Note N1 summarizes these results).

**Figure 1.**
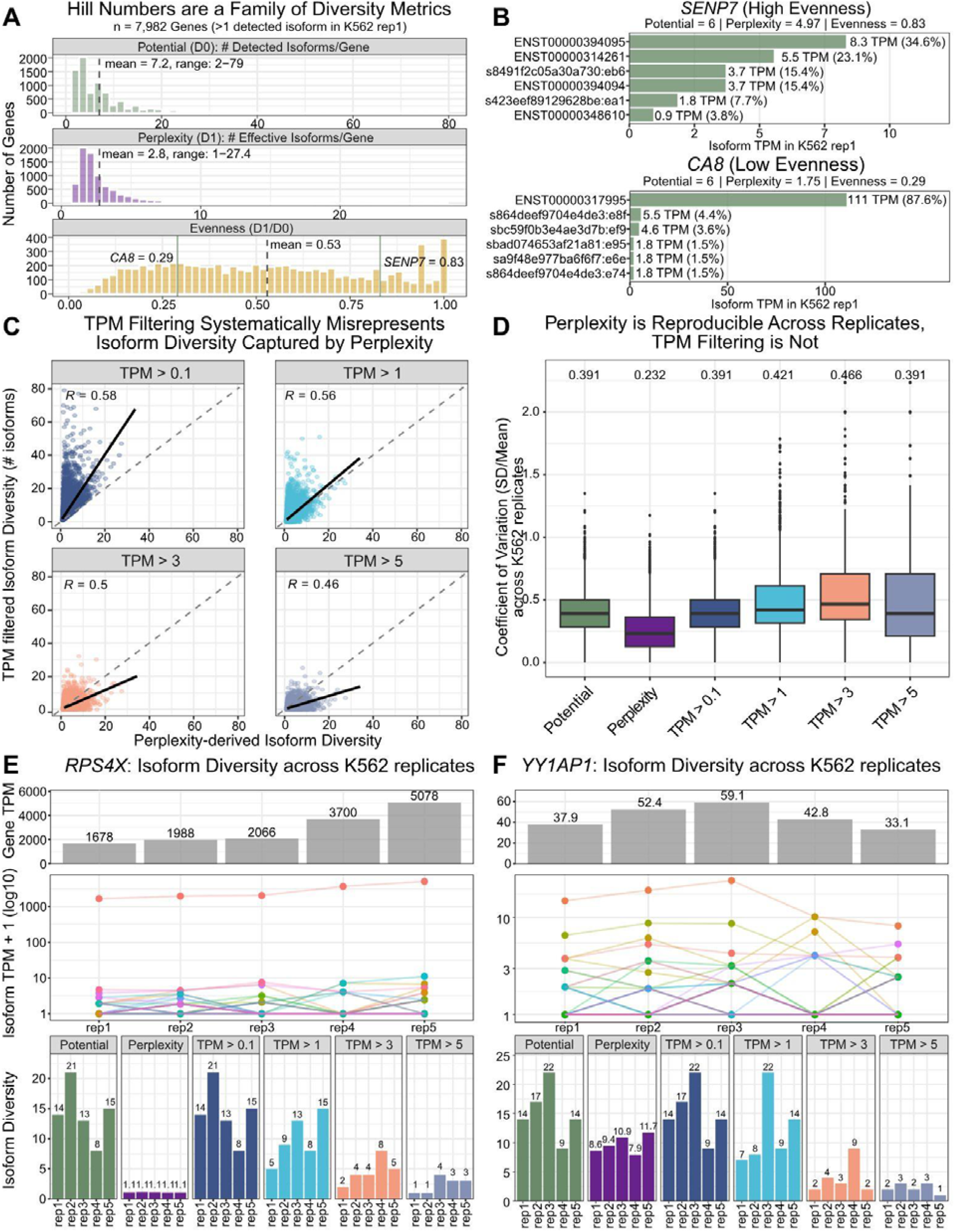
Perplexity is a Principled Metric to Quantify Isoform Diversity. (A) Hill number summaries of potential (D_0_), perplexity (D_1_), and evenness (D_1_/D_0_) per gene in a representative K562 sample. Grey dashed lines mark the mean for each metric. Green lines indicate the evenness of *SENP7* and *CA8*, detailed in B. (B) Illustrative gene examples from K562 showing the isoform abundance distribution for *SENP7* (high evenness) and *CA8* (low evenness). (C) Per-gene comparison of isoform diversity estimated by perplexity versus the number of isoforms remaining after applying a TPM threshold of 0.1, 1, 3, or 5. Pearson’s correlation (*R*) and a linear regression line in black are shown in each panel. The dashed diagonal grey lines represent a slope of 1 where the two estimates would be equal. All correlations are significant (p < 2.2 x 10^-16^). (D) Per-gene coefficient of variation (CV = standard deviation/mean) of isoform diversity estimates across five K562 replicates. Values at the top are the median CVs. Potential and TPM > 0.1 boxplots are identical as a filter of 0.1 keeps all isoforms per gene. (E-F) Isoform diversity metrics across five K562 replicates for RPS4X (E), a low-evenness gene, and YY1AP1 (F), a high-evenness gene, each with many isoforms near typical TPM thresholds. Per replicate gene TPM (top), isoform TPM (middle), and diversity estimates from perplexity and TPM filtering (bottom).

The framework captures isoform diversity at two orders: *potential* (the number of observed isoforms, D_0_) and *perplexity* (the *effective* number of isoforms, D_1_), from which *evenness* (D_1_/D_0_) is derived (Figure 1A). To introduce our framework, we focus on a representative K562 sample. We compare two genes, *SENP7* and *CA8*, each with six detected isoforms (Figure 1B).

Potential (D_0_) marks the upper bound of a gene’s isoform diversity: the total number of detected isoforms per gene. While *SENP7* and *CA8* share a potential of 6, potential alone does not fully capture the isoform profile. Perplexity (D_1_) condenses the abundance distribution of detected isoforms into a single value per gene: the effective number of isoforms, where each isoform is weighted by its relative abundance. For *SENP7* (23.9 TPM), expression is distributed across all D_0_=6 isoforms (0.9-8.3 TPM), and perplexity (4.97) approaches potential, indicating that isoforms contribute meaningfully to diversity. In contrast, *CA8* (126.5 TPM) expresses one dominant isoform at 111 TPM and five isoforms each contribute less than 5%. Perplexity reflects this and collapses to 1.75 effective isoforms, despite the starting potential of 6. Evenness (D_1_/D_0_) quantifies these differences in isoform expression distributions on a scale from 0 to 1, where *SENP7* scores 0.83 (even; 4.97/6) and *CA8* scores 0.29 (uneven; 1.75/6).

The standard approach to estimating isoform diversity is to apply a global expression threshold and count the remaining isoforms per gene. The *SENP7* and *CA8* examples expose fundamental limitations of this approach– that no single threshold is satisfactory for all genes simultaneously, and the choice of threshold systematically misrepresents isoform diversity. At low thresholds (TPM > 0.1), nearly all isoforms are retained, overestimating diversity, especially for low-evenness genes like *CA8*. As thresholds increase to 1, 3, or 5 TPM, estimates improve for low-evenness genes but underestimate diversity for high-evenness genes like *SENP7* (Figure 1C). Perplexity requires no such threshold; it is a mathematically principled diversity metric that adapts to each gene’s unique isoform abundance distribution.

In addition to systematic misrepresentation, threshold-based estimates of isoform diversity are unstable across replicates. Small shifts in TPM can push isoforms above or below a threshold, producing high batch-to-batch variation in diversity measurements. We quantify this across five K562 replicates (Figure 1D). Because perplexity considers the full abundance distribution rather than applying a binary cutoff, it is more robust to small shifts in TPM. Two genes, *RPS4X* and *YY1AP1*, further illustrate this contrast across the five replicates (Figure 1E-F). *RPS4X* is a low evenness gene, expressing a dominant isoform and many low-abundance isoforms that fluctuate near typical TPM thresholds. Perplexity is stable at 1.1 across all five replicates, while TPM-filtered estimates range from 5-15 and 1-4 at a 1 and 5 TPM threshold, respectively (Figure 1E). *YY1AP1* is a high evenness gene with total expression between 33.1 and 59.1 TPM across replicates, but each individual isoform is expressed near typical thresholds.

Perplexity is high, ranging from 7.9-11.7 effective isoforms across replicates compared to 7-22 and 1-3 at 1 and 5 TPM cutoffs, respectively (Figure 1F).

Together, these results demonstrate that perplexity provides a principled alternative to expression-based filtering for measuring isoform diversity. Rather than imposing a binary decision for each isoform, perplexity considers each isoform’s contribution to a gene’s diversity– whether large or small. It adapts across genes that vary in overall expression and isoform complexity, and is robust to variation across replicates.

### Perplexity Reveals the Isoform Diversity Landscape Across 124 ENCODE4 LRS Samples

In the five K562 replicates from Figure 1, we detected 96,545 isoforms across 9,578 genes. To assess isoform diversity on a larger scale, we applied potential and perplexity to all 124 ENCODE4 PacBio LRS datasets across 55 cell and tissue types (Reese et al. 2023). After selecting for genes with at least two protein-coding isoforms, we retained 12,658 genes and all their associated isoforms, including nonsense-mediated decay (NMD), retained intron (RI), and no open reading frame (noORF) biotypes (Supplemental Figure 1B-E, Supplemental Table S1-S2). For each gene, perplexity was calculated from isoform ratios averaged across all samples where the gene is expressed, capturing the isoform diversity landscape across samples.

As expected, the number of detected isoforms increases with each additional tissue, reaching 185,160 across the full dataset (Figure 2A). SQANTI3 classifies 78% and 74% of these isoforms as ‘novel in catalog’ and ‘novel not in catalog’ (NIC and NNC) (Pardo-Palacios, Arzalluz-Luque, et al. 2024) based on GENCODE v46 basic and comprehensive annotations, respectively, consistent with prior LRS analyses (Supplemental Figure 2A) (Monzó et al. 2025; Pardo-Palacios, Wang, et al. 2024; Reese et al. 2023; Glinos et al. 2022). In the full dataset, the average per-gene potential is 14.6 detected isoforms (range: 2-316) and the average per-gene perplexity is 3.4 effective isoforms (range: 1-47.1) (Figure 2B). Similar to Figure 1A, evenness varies widely across genes (mean: 0.4; Figure 2B). This is further demonstrated in a direct comparison of per-gene potential and perplexity (Figure 2C).

**Figure 2.**
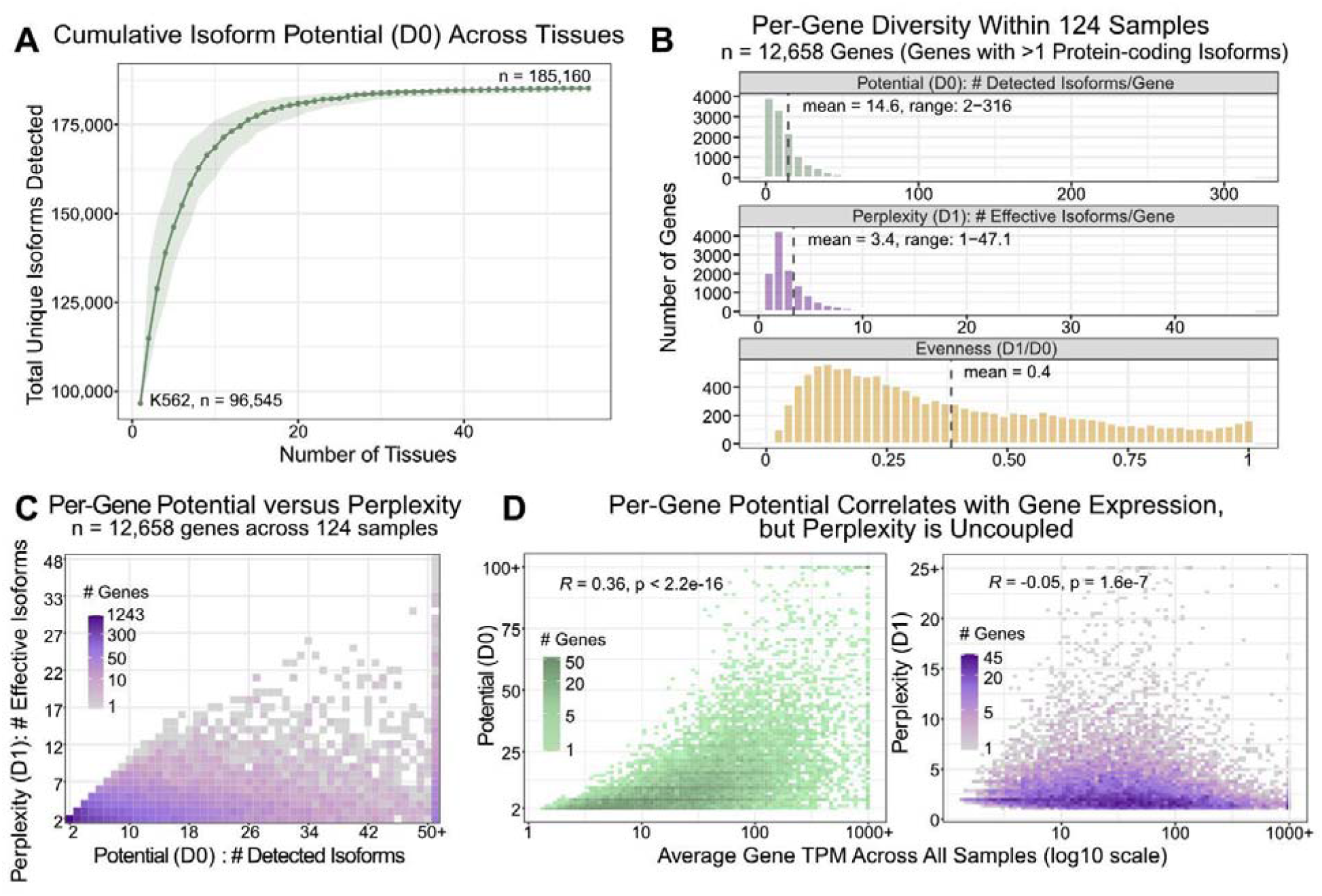
Perplexity Reveals the Isoform Diversity Landscape Across 124 ENCODE4 LRS Samples. (A) Cumulative number of unique isoforms detected as tissues are added, starting with K562. The line and points represent the median across 100 random tissue orderings and the shaded area indicates the 5th-95th percentile range. (B) Hill number summaries of potential (D_0_), perplexity (D_1_), and evenness (D_1_/D_0_) per gene within all 124 ENCODE4 LRS samples. Grey dashed lines mark the mean for each metric. (C) Per-gene potential compared to per-gene perplexity for genes with >1 protein-coding isoforms. Color intensity corresponds to the number of genes at each point. (D) Average gene expression versus per-gene potential (left) and per-gene perplexity (right). Samples with a gene expression of 0 are excluded from the average. Color intensity corresponds to the number of genes at each point. Pearson’s *R* is shown.

Previous studies using ESTs or short read RNA-seq have reported that gene expression and exon count are positively correlated with isoform diversity (Melamud and Moult 2009; Pickrell et al. 2010; Tapial et al. 2017). This relationship is expected: highly expressed genes are more deeply sampled and genes with more exons have more opportunities for variable splice junction usage, both increasing the likelihood of detecting multiple isoforms. Indeed, in the ENCODE4 LRS data, per-gene potential showed a modest positive correlation with gene expression (*R* = 0.36; Figure 2D), and a correlation, albeit weaker, with exon count (*R* = 0.16; Supplemental Figure 2B). In contrast, per-gene perplexity is uncoupled from gene expression, showing only a very weak *negative* correlation (*R* = -0.05; Figure 2D). Perplexity showed a similar association with exon count (*R* = 0.21; Supplemental Figure 2C), but this relationship is likely confounded by technical bias for shorter read lengths–and thus shorter genes.

Overall, expression and exon count explain only 4.5% of the variance in perplexity, compared to 15.8% for potential. This discrepancy suggests that the correlation between potential and expression is primarily driven by technical factors–such as increased detection power in highly expressed genes–rather than reflecting genuine differences in isoform regulatory complexity. In contrast, perplexity’s independence from expression underscores its generalizability across genes regardless of expression level, capturing intrinsic isoform diversity.

### Perplexity to Evaluate Isoform Diversity Across Regulatory Levels

In Figures 1-2, perplexity (D_1_) was calculated using all detected transcripts of protein-coding genes, including protein-coding, NMD, RI, and noORF biotypes. However, perplexity can also be recalculated on subsets of transcripts to capture different layers of gene regulation. For example, excluding non-coding transcripts (i.e., NMD, RI, noORF) and re-scaling expression ratios among the remaining protein-coding transcripts yields *protein-coding (pc) transcript perplexity*, which reflects diversity among translatable transcripts. To focus more specifically on protein-level diversity, transcripts that encode the same open reading frame (ORF) can be collapsed to give *ORF perplexity*, reflecting instead the effective number of distinct protein products produced by each gene (Figure 3A).

**Figure 3.**
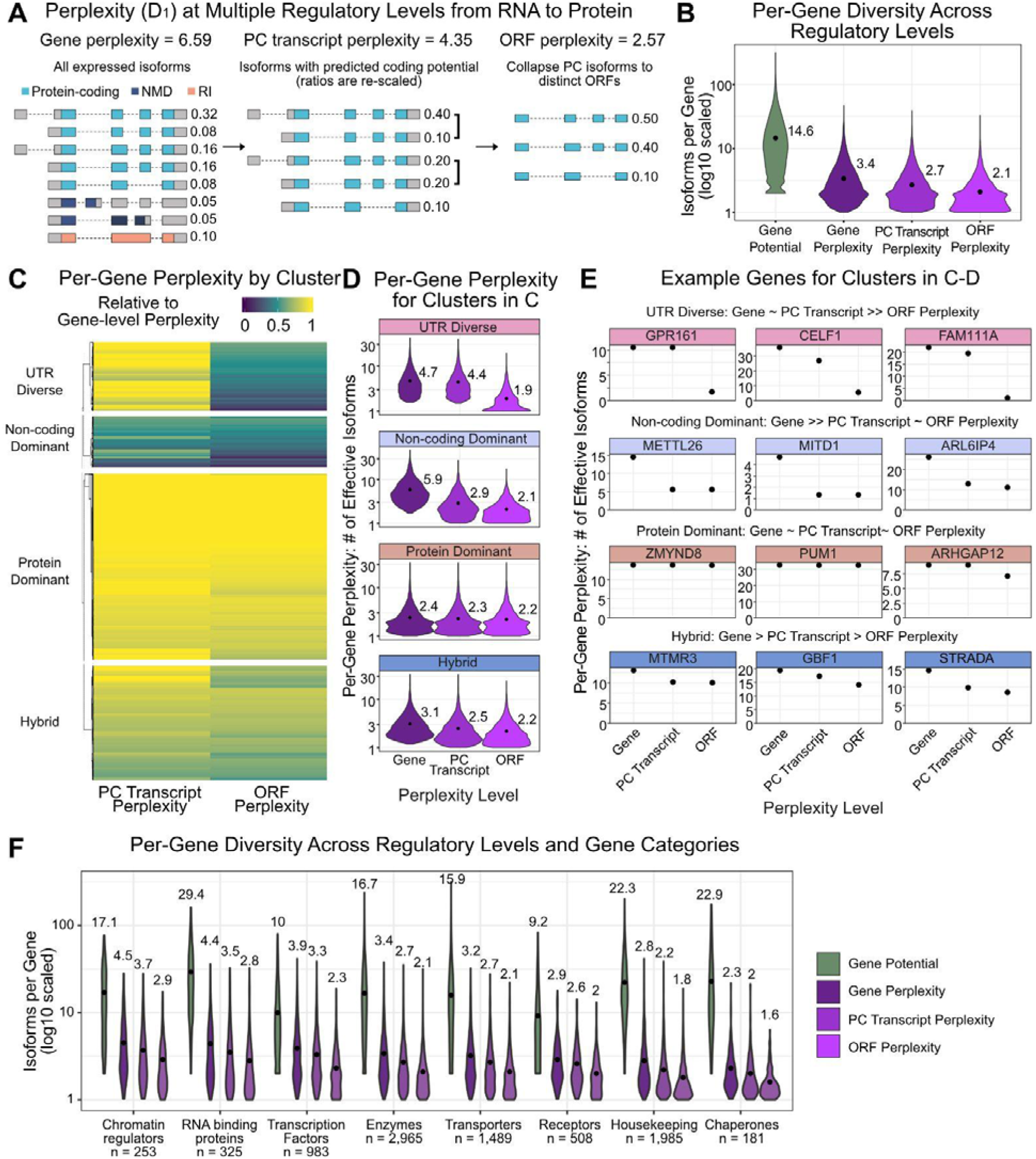
Perplexity to Evaluate Isoform Diversity Across Regulatory Levels. (A) Per-Gene isoform diversity can be evaluated at three regulatory levels: gene, protein-coding (pc) transcript, and ORF. Exon colors indicate isoform biotype (protein-coding, NMD, RI). Expression ratios are given on the right. (B) Per-gene diversity shown as potential and perplexity at three regulatory levels: gene, pc transcript, and ORF (for genes with multiple protein-coding transcripts, n = 12,658). Numbers next to the violins are the average isoforms per gene for each metric. (C) Genes (y-axis) are hierarchically clustered (Euclidean distance) by the relative change in perplexity from gene to pc transcript and ORF levels. Four groups are labelled based on these patterns. (D) Distribution of per-gene perplexity at each regulatory level for the four gene groups in (C). Numbers give the average perplexity. (E) Three example genes from each cluster in C-D are plotted. For each gene, gene-level perplexity is plotted alongside pc transcript and ORF perplexity. The “∼” indicates approximately equal, “>” indicates moderately greater, and “>>” indicates substantially greater. (F) Per-gene potential and perplexity at each regulatory level across functional gene categories. The average is labelled above each violin. The x-axis is ordered by the average ORF perplexity. The number of genes in each category are given on the x-axis. All pairwise comparisons in (B) and within each group in (D) are significant (Wilcoxon signed-rank test, p < 10^-16^).

Among genes with multiple protein-coding transcripts (n = 12,658), isoform diversity decreases at each regulatory level: from an average gene perplexity (all transcripts) of 3.4 to a perplexity of 2.7 for pc transcripts and 2.1 for ORFs (Figure 3B). The final value suggests that, on average, these genes produce approximately two distinct proteins that are reliably expressed across samples. Across genes, however, the degree of change at each level varies greatly, and we observe four broad patterns based on how perplexity changes across levels (Figure 3C-E). *UTR Diverse* genes show similar gene and pc transcript perplexity but low ORF perplexity, indicating that most RNA isoforms are protein-coding and share the same ORF, differing primarily in untranslated regions (top row). *Non-coding Dominant* genes exhibit a sharp decrease from gene to pc transcript perplexity, suggesting that most of their RNA diversity comes from non-coding transcripts, such as NMD, RI, and noORF (2nd row). *Protein Dominant* genes show minimal change across all three levels, indicating that most of their isoforms are protein-coding and produce distinct proteins (3rd row). *Hybrid* genes fall between these extremes with moderate reductions in perplexity across levels, suggesting a mix of functional and redundant isoform diversity (4th row). Despite these distinct patterns, all four groups converge on a similar average ORF perplexity between 1.9-2.2 (Figure 3D).

Patterns of perplexity across levels also vary by functional gene categories (Figure 3F). Genes encoding regulatory proteins, such as chromatin regulators, RNA binding proteins, and transcription factors show the highest perplexity at all levels (Lambourne et al. 2024; Talavera et al. 2009), with ORF perplexities of 2.9, 2.8, and 2.3, respectively. In contrast, housekeeping genes and chaperones have high average gene potential but the lowest ORF perplexities, likely reflecting pervasive low-level transcription rather than functional diversification. These trends highlight that even though housekeeping genes may express a far higher number of distinct transcripts overall, regulatory gene classes exhibit greater proteomic diversity.

### ORF Expression Breadth and Variability Across Tissues

In Figures 2-3, perplexity was calculated using isoform usage ratios averaged across all ENCODE samples, providing a global estimate of isoform diversity. However, this aggregate view obscures tissue-level heterogeneity, i.e., whether multiple ORFs are co-expressed within individual samples or partitioned across tissues. We therefore calculated perplexity on a per-sample basis within all 124 samples from 20 organ types (Figure 4A; Supplemental Table S3). Within each sample, we classified ORFs as effective or ineffective by rounding each gene’s ORF perplexity to the nearest integer and designating that number of highest-expressed isoforms as effective (Figure 4B). From this framework, we derived two complementary continuous measures of tissue-specificity per-ORF: expression breadth and variability.

**Figure 4.**
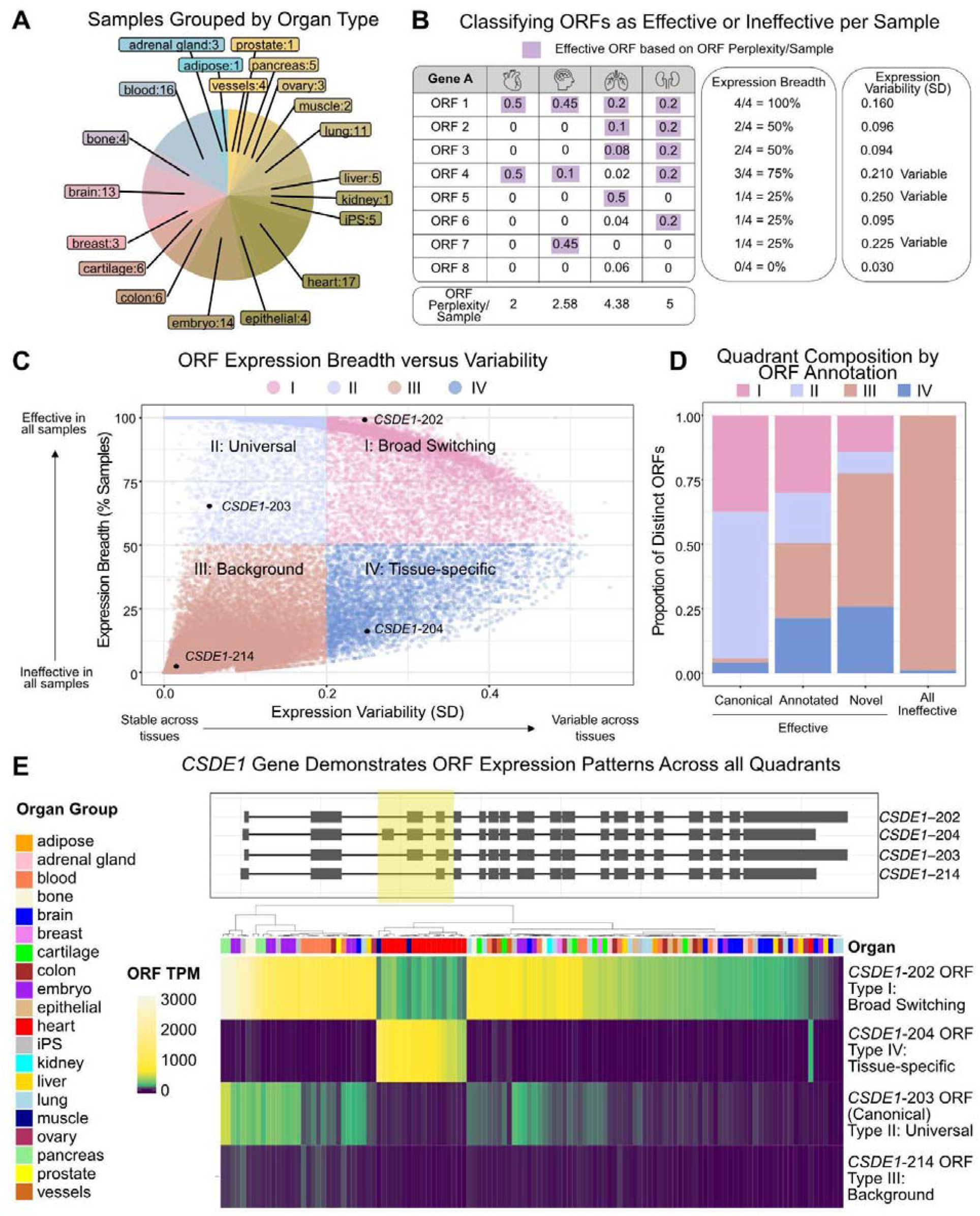
ORF Expression Breadth and Variability Across Tissues. (A) 124 ENCODE4 PacBio LRS samples grouped by organ type (Supplemental Table S3). The number of samples per organ type is shown. (B) ORF perplexity calculated per sample to give *Expression Breadth* and *Expression Variability*. ORFs in purple boxes are those classified as effective in each sample. ORF ratios are only included for samples where the gene is expressed. (C) ORFs are plotted by expression variability (SD; x-axis) versus expression breadth (% samples effective; y-axis) and grouped into quadrants. The *CSDE1* gene has ORFs across all four quadrants–one ORF from each quadrant is marked. (D) Quadrant proportion of ORFs classified as effective in the aggregate analysis, stratified by mutually exclusive annotation (canonical, annotated, novel) alongside all aggregate-ineffective ORFs. (E) Top: Structure of the four *CSDE1* ORFs marked in C. The exons that differ between the isoforms are highlighted in yellow. Bottom: Sample-specific TPM of these four *CSDE1* ORFs. Organ types are labelled at the top. The canonical ORF (203) is labelled.

Expression breadth quantifies how consistently each ORF is expressed, defined as the percentage of samples in which an ORF is classified as effective. For example, ORF 1 is effective in all 4 samples (100% expression breadth), while ORFs 5 and 7 are only effective in a single sample (25% expression breadth, Figure 4B). As a complementary measure, expression variability—the standard deviation of ORF usage ratios across samples— captures how much an ORFs relative expression fluctuates across samples. For instance, an ORF that shifts from a usage ratio of 0.1 to 0.9 is more variable, and potentially more tissue-specific, than one that remains consistently between 0.8 and 0.9.

Tissue-specificity is continuous, although often treated as binary. Expression breadth and variability capture this continuity in two dimensions (Figure 4C). To facilitate interpretation of patterns across this continuous space, we overlaid descriptive quadrant boundaries at 50% expression breadth and 0.2 variability. These boundaries are heuristic, not firm thresholds– continuous expression breadth and variability for all ORFs are provided in Supplemental Table S1. Quadrants I (*Broad Switching*) and II (*Universal*) include ORFs with high expression breadth that differ in variability. Together, they account for the vast majority of canonical ORFs (GENCODE v46 Ensemble canonical), underscoring their widespread expression signatures (37% and 57%, respectively) (Figure 4D; Supplemental Figure 3A). Quadrant III (*Background*) is defined by ORFs with both low breadth and low variability and contains the majority of ineffective ORFs (99%). Quadrant IV, (*Tissue-specific*) captures ORFs with low breadth but high variability–potential markers of context-specific usage. We find that a large proportion of novel and annotated effective ORFs fall into Quadrant IV, 26% and 21%, respectively. In contrast, canonical (4%) and ineffective (1%) ORFs are rarely classified as tissue-specific (Figure 4D; Supplemental Figure 3A). Therefore, tissue-restricted isoform diversity disproportionately arises from non-canonical and novel ORFs.

To examine whether ORFs classified as tissue-specific (Quadrant IV; n = 3,973) exhibit coherent tissue-restricted usage patterns, we hierarchically clustered usage ratios per sample for these ORFs, resulting in ten clusters. We observed the most pronounced tissue-resolved clusters emerge for brain, heart, and embryonic tissues, which have the greatest sample representation in the ENCODE4 data (Supplemental Figure 3B; Supplemental Table S4).

Tissue-level pattern resolution is limited for tissues with only one or two samples. In addition, receptors, transcription factors, and chromatin regulators have the highest proportion of tissue-specific ORFs, while housekeeping genes have the lowest (Supplemental Figure 3C).

The *CSDE1* gene, an RNA-binding and translation regulator, illustrates how distinct isoforms from the same gene exhibit different expression patterns, with at least one ORF in each quadrant (Figure 4C,E). *CSDE1-202* is broadly expressed except in heart and muscle, where *CSDE1-204* is dominant (Figure 4E). Both ORFs show high variability across samples, but differ in tissue breadth so that *CSDE1-202* falls into Quadrant I (broad switching) while *CSDE1-204* is in Quadrant IV (tissue-specific). These isoforms differ by a cassette exon included in 204 and skipped in 202 (Figure 4E). In comparison, *CSDE1-203*, the canonical ORF, shows moderate, but consistent expression across samples and falls into Quadrant II (Universal), while *CSDE1-214*, with minimal and uniform expression, belongs to Quadrant III (Background) (Figure 4E).

The cardiac/muscle-specific alternative exon in *CSDE1-204* was identified previously (Salomonis et al. 2009) but isoform-specific functions of *CSDE1* remain poorly characterized. Notably, this gene has been implicated in cardiovascular disease, neurodevelopmental and psychiatric disorders, and cancer (Ciocia et al. 2024), highlighting the potential for isoform-specific disease relevance.

## Discussion

The way we define transcriptome composition–the full set of RNA molecules generated by the genome–is a foundational decision that influences nearly every layer of downstream biological interpretation. It shapes our understanding of gene output, guides hypothesis generation, informs the interpretation of disease-associated variants, and delineates the parameter space for system-level modeling. For this critical step, the field currently relies on user-defined thresholds that involve an ad hoc mix of TPM cutoffs, cross-sample expression support, or isoform abundance ratios, resulting in widely divergent estimates of transcriptome complexity. In this study, we introduced perplexity as a principled alternative to quantify transcriptome diversity.

With perplexity, all detected isoforms, regardless of expression level, contribute proportionally to a gene’s diversity, rather than being excluded as presumed transcriptional and splicing noise (Melamud and Moult 2009; Wan and Larson 2018). This approach is consistent with the observation that molecular diversity is already far more biologically constrained than the theoretical space of all possible isoforms. For example, the *TTN* gene contains 363 exons, which could theoretically produce 4.7 x 10^108^ isoforms through exon skipping alone, without considering alternative start and ends or other splicing events. Yet only 14 protein-coding isoforms of *TTN* are annotated in GENCODE, highlighting the remarkable specificity of the transcriptional and splicing machinery (Deveson et al. 2018).

Perplexity is generalizable across regulatory levels: gene, transcript, and protein. This captures both the extent and source of a gene’s diversity at different levels of granularity–whether arising from UTRs, non-coding transcripts, or distinct ORFs. These sources of variation suggest distinct regulatory mechanisms, from dosage control via NMD or translational regulation to the production of distinct protein products (Fair et al. 2024; Nilsen and Graveley 2010; Rogalska et al. 2023). When applied across all samples to protein-coding genes, perplexity revealed that genes express an average of 3.4 transcript isoforms and 2.1 protein isoforms, offering a data-driven estimate of isoform diversity that does not depend on arbitrary filtering choices.

Perplexity is first and foremost a diversity metric, as demonstrated in Figures 1-3. In Figure 4, we extend its use to classify ORFs as effective or ineffective on a per-sample basis. The properties that make perplexity a robust diversity metric, adaptability to gene-specific expression and complexity and reproducibility across replicates (Figure 1), also support its use as a principled threshold. Unlike fixed absolute expression thresholds (TPM), which favor highly expressed genes and penalize complex ones, perplexity applies an adaptable threshold per gene. Minor variation in expression across samples has a negligible effect on perplexity and thus on the resulting classification. We acknowledge that any threshold–perplexity or TPM based–has limitations. In the absence of ground truth distinguishing biologically relevant isoforms from transcriptional or splicing noise, no threshold can be objectively justified and any threshold will retain some noisy isoforms and remove some functional ones. The advantage of perplexity is that it is systematic and rooted in the Hill number framework rather than arbitrary. We also note that some research questions may necessitate absolute expression thresholds, such as in isoform-specific therapeutic targeting, where an isoform must be expressed above a minimum level to be effectively targeted.

While perplexity offers an improved framework to quantify isoform diversity, it remains an expression-based metric. As such, it operates under the assumption that relatively abundant RNA isoforms are more likely to be biologically consequential. However, RNA abundance represents only one dimension of biological consequence, and its relationship to downstream protein output is imperfect. RNA and protein abundance are only moderately correlated (Gry et al. 2009; Jiang et al. 2020; Schwanhäusser et al. 2011). Low-abundance isoforms can be efficiently translated or result in highly stable proteins (Weatheritt et al. 2016; Blencowe 2006), 5’UTR differences can affect translation efficiency, and N-terminal amino acids and internal isoform-specific sequences can influence protein stability (Varshavsky 2011). Therefore, any isoform expression-based proxy for functional impact should be paired with direct detection of protein isoforms, a nascent area of study (Miller et al. 2022; Mehlferber et al. 2022; Sinitcyn et al. 2023; Kedan et al. 2025), as well as functional validation.

Despite the scale of the ENCODE4 LRS data, the isoform catalog analyzed here almost certainly underestimates isoform diversity. Many cell types have not been sequenced meaning current catalogs likely lack many tissue-specific isoforms. In addition, LRS faces two main technical limitations (Reese et al. 2023; Pardo-Palacios, Wang, et al. 2024; Calvo-Roitberg et al. 2024; Mikheenko et al. 2022). First, incomplete reverse transcription (RT) processivity and PCR amplification bias during library preparation lead to underrepresentation of long transcripts (>12 kb) (Reese et al. 2023). Second, TSS annotation is complicated by read truncation from RNA degradation, internal priming, or incomplete RT (Pardo-Palacios, Arzalluz-Luque, et al. 2024).

To minimize overcalling TSS diversity, we collapsed reads to isoforms without requiring identical 5’ ends, with the consequence of underestimating 5’UTR variation in our study. Overcoming these technical barriers remains an active area of development with improved RT enzymes, direct RNA-sequencing, 5’ cap-capture methods, and validation with orthogonal datasets (CAGE for TSS verification) (Ogami et al. 2023; Pardo-Palacios, Arzalluz-Luque, et al. 2024; Garalde et al. 2018).

We acknowledge that our perplexity estimates for lowly expressed genes will be slightly downward biased (i.e., we slightly underestimate perplexity) because of limited coverage (Miller and Miller 1955). We considered applying existing bias correction approaches, but these have their own limitations: Miller-Madow bias correction requires knowing the true number of possible isoforms (Miller and Miller 1955) and the Nemenman-Shafee-Bialek estimator is computationally expensive to calculate, requiring numerical integration or Monte Carlo (Nemenman et al. 2001). Given the high coverage of the datasets analyzed here, the bias in our estimates will be minimal. This intuition is supported by the slight negative correlation between expression and perplexity in Figure 2D: substantial downward bias for more lowly expressed genes would, in contrast, result in a *positive* correlation.

## Conclusions

Perplexity provides a principled and interpretable measure of isoform diversity across diverse genes, regulatory levels, and cell types. It considers all high-confidence transcript structures– including low-abundance ones–to contribute proportionally to a gene’s diversity. It is easy to compute, straightforward to interpret, and immediately deployable in existing workflows. Beyond LRS transcriptomics, the perplexity framework is applicable to any omics dataset where molecular diversity can be quantified from relative abundances. Our analysis pipeline is available on GitHub, along with IsoPlex, a Python library for calculating perplexity from LRS data.

## Methods

### Customized long read RNA-sequencing analysis pipeline

Our full LRS pipeline is available as a Bash script on GitHub (https://github.com/daklab/Perplexity_Isoform_MasterTable) and is summarized in Supplemental Figure 1A. The pipeline takes as input raw FASTQ files and a sample-to-tissue mapping file.

#### Download LRS Data from ENCODE4

Data was downloaded from ENCODE4 on September 30th, 2024, in FASTQ format. The dataset was filtered for unperturbed, *Homo sapiens*, long read RNA-seq on PacBio Sequel II and Sequel platforms. The polyadenylated mRNA sequences were assembled on GRCh38. Based on these filters, 124 samples were used in this study.

#### Read Alignment with Minimap2

Raw reads were aligned with Minimap2 v2.17 (Li 2018) using recommended parameters for PacBio long-read sequences: -ax splice:hq -uf. Reads with MAPQ smaller than 60 were filtered out, and only primary aligned reads were kept. BAM output files were converted to BED and PSL files using helper scripts from FLAIR v1.6.1 (Tang et al. 2020): bam2Bed12.py and bed_to_psl.py.

#### Collapsing Reads to Isoforms

We wrote a custom script, available on GitHub (sp_collapse_isoforms.py), that utilizes the logic of FLAIR collapse (Tang et al. 2020) to collapse full length reads to isoforms based on shared junction coordinates. Only internal junctions were considered in this step–exact read start and ends were not considered. TES and TSS of an isoform were defined by the supporting read with the maximum UTR length. Single-exon reads were filtered out. Resulting isoforms were output as PSL and GTF files.

#### Removal of Technical Artifacts

Before applying the Hill number framework, it is essential to remove technical artifacts, especially abundant artifacts such as internal priming, as this can inflate perplexity. We applied stringent filtering prior to any diversity calculations.

Internal priming and fragment artifacts can be highly abundant and, if retained, would inflate diversity estimates. To remove internal priming artifacts, we required each read to overlap a GENCODE v46 basic annotated last exon (transcription end sites; TES). Next, to remove non-full length reads which included fragments, fragments with retained introns, and reads that aligned fully within the 3’UTR, we removed reads that did not overlap an annotated first exon (transcription start sites; TSS). The number of reads removed at each step is shown in Supplemental Figure 1A.

To remove remaining technical artifacts that are more challenging to filter out bioinformatically (RT template switching and sequencing errors at splice junctions), we applied a minimal filter that required 10 supporting reads per isoform across all 124 samples. At this step, 11,551,729 reads (∼5% of raw reads) belonging to 7,894,625 collapsed “isoforms” are removed–81% of these isoforms have only one supporting read across all 124 samples. Without this filter, these artifacts would drastically inflate potential but have negligible effects on perplexity, as single-read isoforms contribute minimally to the abundance distribution.

#### Overlap with GENCODE v46 Annotations

Isoforms with all internal junctions matching with annotated transcripts from GENCODE v46 were assigned with annotated transcript IDs. If more than one annotated transcript matched a collapsed isoform–for two annotated transcripts that only differ in UTR lengths–their transcript IDs were appended with a semicolon. For all isoforms, including the novel ones, Ensembl gene IDs from GENCODE v46 were assigned to isoforms using a FLAIR helper script (identify_gene_isoform.py). Unannotated genes are assigned with chromosome loci for their gene ID.

A Python v3.11.5 hash function (sha256) was used to generate reproducible novel isoform ids based on exon start and end coordinates (sp_collapse_isoforms.py).

#### Identifying Retained Introns with SQANTI3

We ran SQANTI3 v5.1.1 QC (Pardo-Palacios, Arzalluz-Luque, et al. 2024) to identify isoforms with retained introns (RI). All isoforms categorized as “intron_retention” were marked as RI for further analysis.

#### Calling Open Reading Frames (ORFs) with CPAT

CPAT v3.0.4 was used to predict open reading frames (ORFs) in the remaining isoforms (Wang et al. 2013). To run CPAT, the PSL file was converted to FASTA file format using FLAIR helper script (Tang et al. 2020): psl_to_sequence.py. A prebuilt human logit model and hexamer table were downloaded from CPAT and were used to predict ORFs. A minimum ORF length of 90 nucleotides, 30 amino acids, was required and the top scoring ORF was selected per isoform. We used the faTrans utility v377 from the UCSC Genome Browser tools to translate DNA sequences into amino acid sequences (Karolchik et al. 2004). Isoforms with a predicted ORF were labelled as protein-coding and isoforms with no ORF predicted were labelled as ‘noORF’.

At this step, ORFs were overlapped with GENCODE v46 basic annotations. For annotated ORFs, 94% of CPAT ORF predictions matched the annotated ORF. For the isoforms that CPAT got wrong, we used the annotated ORF. In addition, if a gene is not annotated as “protein_coding” in GENCODE v46 basic, we classify all isoforms pertaining to that gene as “noORF” or “RI”. These isoforms get filtered out in Supplemental Figure S1B, since they have protein-coding isoforms equal to 0, and are not considered throughout the paper. Importantly, this step removes pseudogene alignments.

#### Custom NMD Prediction

A custom script (filter_NMD.py) was used to identify isoforms predicted to undergo nonsense-mediated decay (NMD). The script processes all isoforms with predicted ORFs and classifies an isoform as NMD if it contains a stop codon located upstream of the final splice junction–a premature termination codon (PTC). Additionally, any isoform already annotated as NMD in GENCODE v46 basic were classified as NMD here. Annotated isoforms with GENCODEv46 protein-coding annotation that were predicted to have a PTC were rescued and marked as protein-coding.

#### Filtering for Protein-coding Diversity

Genes with 0 or 1 protein-coding isoforms were filtered out for our further analysis since we focus on protein-coding genes with isoform diversity. Plots in Supplemental Figure 1B-C summarize these isoforms. These isoforms are still contained in the master table.

#### Threshold-based Isoform Diversity Estimates

To compare perplexity against conventional approaches, we applied binary TPM thresholds of 0.1, 1, 3, and 5 to each of five K562 replicates from the ENCODE4 data. Per-gene isoform diversity was estimated by counting the number of isoforms remaining after applying each threshold.

### Master Table Metrics

The master table provides gene, transcript, and ORF level metrics for all 199,406 detected transcripts across 19,226 genes in this study (Supplemental Table S1). Column descriptions are organized and discussed below.

#### Gene, Transcript, and ORF Identifiers

*transcript_id, transcript_name, gene_id, gene_name*: For annotated isoforms, Ensembl (GENCODE v46) gene and transcripts ids are included. For novel isoforms, gene IDs are assigned based on genomic loci using the FLAIR helper script (identify_gene_isoform.py) (Tang et al. 2020). Novel transcript IDs were generated by hashing internal splice junction coordinates using Python’s hashlib library (shake_256). Junction start and end positions were separately hashed to produce a deterministic ID in the format sHASH:eHASH. This ensures that novel isoforms with identical splice junctions receive the same ID regardless of user, time, and environment, enabling cross-referencing across studies. *ORF_id*: Transcript_ids from isoforms encoding identical amino acid sequences are collapsed to a single ORF ID, separated by a semicolon.

#### Expression Metrics

*gene_avg_tpm:* Average gene TPM across ENCODE4 samples in which the gene is expressed. *gene_n_expressed_samples:* Number of samples in which the gene is expressed (max = 124). *transcript_avg_tpm:* Average transcript TPM across all samples in which the gene is expressed. *transcript_ratio:* Transcript expression ratio within its gene averaged across all samples. This is used as the input into gene perplexity calculations. *ORF_ratio:* ORF expression ratio within its gene averaged across all samples.

#### Gene-level Diversity Metrics

*gene_potential:* Total number of detected isoforms per gene across all 124 samples, including protein-coding, NMD, RI, and noORF biotypes. *gene_perplexity:* Effective number of isoforms per gene calculated as **2^*H*^**, where

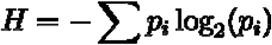

is Shannon entropy and ***p*_*i*_** is a probability distribution of isoform *i* proportions within a gene. For this specific metric, ***p*_*i*_** was computed based on the average isoform ratios across all 124 ENCODE samples. *ptc_potential*: Total number of detected protein-coding isoforms per gene. *ptc_perplexity*: Effective number of protein-coding isoforms per gene, calculated on re-scaled expression ratios among protein coding isoforms only. *ORF_potential*: Total number of distinct ORFs per gene. *ORF_perplexity:* Effective number of distinct ORFs per gene, calculated on ORF-level ratios where isoform ratios sharing the same ORF are summed.

#### Per-sample ORF Metrics

*ORF_n_samples_effective:* Number of samples in which the ORF is classified as effective. *ORF_expression_breadth:* Percentage of samples in which the ORF is classified as effective (ORF_n_samples_effective/gene_n_expressed_samples).

*ORF_expression_variability:* Standard deviation of ORF ratios across samples where the gene is expressed. *ORF_quadrant:* Quadrant assignment (I–IV) based on expression breadth and variability (Figure 4C).

#### Classifications

*transcript_biotype:* Protein-coding, nonsense-mediated decay (NMD), retained intron (RI) or no open reading frame (noORF) as classified in our custom analysis pipeline. *transcript_n_exons:* Number of exons per transcript. *canonical_gencode_v46*: TRUE if the transcript’s splice junction chain matches a GENCODE v46 transcript tagged as ‘Ensemble Canonical’. *annotated_gencode_v46*: TRUE if the transcript’s splice junction chain matches any transcript in GENCODE v46 basic annotation. *transcript_effective*: TRUE if the isoform is classified as effective based on gene perplexity rounded to the nearest integer. *ORF_effective*: TRUE if the ORF is classified as effective based on ORF perplexity rounded to the nearest integer. *gene_protein_category:* Functional gene category based on Protein Atlas (https://www.proteinatlas.org) (Uhlén et al. 2015), ENCODE for RNA-binding proteins (Van Nostrand et al. 2020), and Housekeeping Atlas (Hounkpe et al. 2021). Categories include RNA-binding proteins, transcription factors, enzymes, receptors, transporters, chaperones, chromatin regulators, actin-binding proteins, and housekeeping genes. Genes without a category here were labelled as ‘Other’; genes in more than one category include all labels.

## Declarations

### Availability of data and materials

No new sequencing data was generated in this study. All PacBio long read RNA-sequencing data processed in this study was previously published through ENCODE4 and is publicly available at: https://www.encodeproject.org/. IsoPlex Python library to calculate perplexity metrics is on github (https://github.com/fairliereese/isoplex) with additional documentation at https://fairliereese.github.io/isoplex/. The bash script containing the full analysis pipeline is available on github (https://github.com/daklab/Perplexity_Isoform_MasterTable). The master table generated as a part of this study is included as Supplemental Table S1.

### Competing Interests

G.M.S. is on the scientific advisory board of Quantum-Si Incorporated and holds stock in Quantum-Si Incorporated.

### Funding

M.D.S. was supported by NIH/NIGMS (F32GM142213). S.H.P. was partially supported by a Columbia University Fu Foundation School of Engineering and Applied Science Bonomi Scholarship and Columbia University Scholars Program Summer Enhancement Fellowship. G.M.S. is supported by NIH/NCI (R33CA281919). D.A.K. is supported by Columbia University and NYGC startup funds, the MacMillan Center for the Study of the Non-Coding Cancer Genome, NIH/NCI (R21CA272345), and NSF CAREER DBI2146398. The content is solely the responsibility of the authors and does not necessarily represent the official views of the National Institutes of Health.

### Author Contributions

M.D.S. and D.A.K. conceived the study. M.D.S., S.H.P, and J.S performed analyses. F.R. and M.D.S. developed IsoPlex. M.D.S., S.H.P, G.M.S., and D.A.K. wrote the paper with input from co-authors. M.D.S, G.M.S., and D.A.K. supervised the study.

## Notes

### Summary of Updates

This revision incorporates peer review feedback. We have added background on Hill numbers, a new Figure 1 comparing perplexity against TPM filtering at four thresholds, and a robustness analysis across K562 replicates. We include the IsoPlex Python library for calculating perplexity in long-read sequencing data.

