## Supplemental Note for "Perplexity as a Metric for Isoform Diversity in the Human Transcriptome"

There is a long history in ecology of using what we call *perplexity*, under the name *Shannon diversity* or the *Hill number of order 1*, as a measure of species diversity/biodiversity. We prefer the term perplexity since it is unambiguously the exponent of the Shannon entropy (in nats), rather than the Shannon entropy itself, whereas "Shannon diversity" has been used to denote both of these quantities.

Here, we summarize the theoretical characterization of Hill numbers, and the special case of perplexity, developed in Leinster 2021. Firstly, we specify some desiderata that we would like a diversity metric to have. Then we show that the Hill numbers are the only diversity metrics that satisfy all these desiderata. Finally, we note some properties of Hill numbers.

First we need some basic definitions.

**Definition 1** (Simplex).

$$\Delta_n := \left\{ p \in (0,1]^n \;\middle|\; \sum_{i=1}^{n} p_i = 1 \right\}$$

**Definition 2** (Uniform distribution over $n$ categories). $u_n = (1/n, \ldots, 1/n)$.

We consider functions $D$ that map from the simplex to positive reals (technically we consider the sequence of functions indexed by $n$). What properties might we want $D$ to have to be a meaningful measure of *diversity*?

- *Symmetry*: $D(p)$ is invariant under permutations of $p$.
- *Absence-invariance*: Adding zero-probability categories does not change $D$.
- *Continuity in positive probabilities*: $D$ is continuous on $\Delta_n$.
- *Normalization*: $D(u_1) = 1$, i.e. if there is only one category then the diversity is 1.

For two more subtle desiderata we need another definition.

**Definition 3** (Product of distributions). *Let $w \in \Delta_n$ and $p \in \Delta_k$. The product distribution $w \otimes p \in \Delta_{nk}$ is defined by,*

$$(w \otimes p)_{i,j} = w_i \, p_j, \qquad 1 \le i \le n,\ 1 \le j \le k,$$

*viewed as a vector in $\Delta_{nk}$ by flattening the index pair $(i,j)$ into a single index.*

**Definition 4** (Replication principle). *$D$ satisfies the replication principle if*

$$D(u_n \otimes p) = n\, D(p), \qquad n, k \ge 1,\ p \in \Delta_k.$$

In ecology the replication principle is motivated by imagining $n$ (equally-sized) islands each with a unique set of $k$ species, and each island having the same $p$. In our context, the replication principle says that, considered together, $n$ genes each with the same distribution of isoform usage $p$ should have diversity $nD(p)$.

**Definition 5** (Modular-monotonicity). *$D$ is* modular-monotone *if*

$$D(p^i) \le D(\tilde{p}^i) \quad \forall i \in \{1, \ldots, n\} \quad \Rightarrow \quad D\big(w \otimes (p^1, \ldots, p^n)\big) \le D\big(w \otimes (\tilde{p}^1, \ldots, \tilde{p}^n)\big),$$

*for all $w \in \Delta_n$, $p^i \in \Delta_{k_i}$, and $\tilde{p}^i \in \Delta_{\tilde{k}_i}$.*

Consider $n$ genes. Gene $i$ has $k_i$ isoforms with isoform usage vector $p^i$ in condition 1 and $\tilde{p}^i$ in condition 2. If the diversity for every gene in condition 1 is less than or equal to that in condition 2, then the combined diversity across genes (weighted by their relative expression $w$ which is assumed the same in both conditions) in condition 1 should also be less than or equal to that in condition 2.

**Definition 6** (Hill numbers). *For $q \in [-\infty, \infty]$, the Hill number of order $q$ is,*

$$D_q(p) := \begin{cases} (\sum_{i=1}^n p_i^q)^{\frac{1}{1-q}}, & q \neq 1, \\ 2^{H(p)}, & q = 1. \end{cases}$$

*where $H(p) = -\sum_{i=1}^n p_i \log_2 p_i$ is the Shannon entropy in bits.*

Perplexity is $D_1$ and what we call potential is $D_0$, the number of nonzero categories (expressed isoforms). General Hill numbers are the exponential of Rényi entropies, which generalize Shannon entropy. One of the many advantages of working with Hill numbers/perplexity rather than entropy itself is it unnecessary to specify the base (i.e., whether the units are bits or nats).

We can now give (a rephrased version of) Theorem 7.4.3 of Leinster 2021:

**Theorem 1** (Characterization of Hill numbers). *$D$ is symmetric, absence-invariant, continuous in positive probabilities, normalized, modular-monotone, and satisfies the replication principle if and only if $D$ is a Hill number.*

If we additionally add the intuitive desideratum that the maximal diversity is attained for $u_n$, then this constrains $q$ to be non-negative (Lemma 7.4.14 of Leinster 2021). Any Hill number with $q \geq 0$ satisfies our desiderata, not just just perplexity ($q = 1$). Perplexity is unique in equally weighting the contribution of every observation (every read in our setting).

We note two properties of Hill numbers (see Lemma 7.4.7 and Equation 4.25 in Leinster 2021):

**Lemma 1** (Hill numbers are multiplicative). *If $D$ is a Hill number and $w \in \Delta_n$ and $p \in \Delta_k$,*

$$D(w \otimes p) = D(w)D(p)$$

Lemma 1 implies the replication principle (but not vice versa without the other desiderata).

Crucial to our interpretation of perplexity is the following:

**Definition 7** (Effective numbers). *$D$ is an* effective number *if*

$$D(u_n) = n \qquad \text{for all } n \geq 1.$$

**Lemma 2** (Hill numbers are effective numbers). *The Hill number $D_q$ is an effective number for any $q$.*
