## Supplemental Figures for "Perplexity as a Metric for Isoform Diversity in the Human Transcriptome"

#### Supplementary Information

##### Supplemental Tables

**Supplemental Table S1: Master Table of Isoforms Detected in ENCODE4 LRS Datasets.** Rows are all 199,406 detected transcripts (19,226 genes) in ENCODE4 PacBio LRS datasets. Columns are summary metrics described in methods. “NA” in a column means that metric was not calculated for that transcript–either because it did not apply or the isoform had already been filtered out.

**Supplemental Table S2: Table of Isoform TPMs Across 124 ENCODE4 LRS Samples.** Rows are all 199,406 detected transcripts (19,226 genes) in ENCODE4 PacBio LRS datasets. Columns are transcripts per million (TPM) across all 124 samples.

**Supplemental Table S3: Organ Type Classifications of ENCODE4 Samples.** The 124 ENCODE4 datasets were aggregated into organ categories based on cell type for Figure 3. Organ groups are mostly similar to (Reese et al. 2023) except for 22 samples defined as ‘in-vitro differentiated cell lines” in Reese et al.–we further classified these here according to more specific organ types.

**Supplemental Table S4: Table of Tissue-specific Quadrant IV ORFs from Figure 4C.** Rows are the 3,973 ORFs from Quadrant IV in Figure 4C. ORFs were hierarchically clustered by Euclidean distance of their ORF usage ratios. The cluster number in the table (1-10) matched the cluster numbers to the right of the heatmap in Supplemental 3B. The continuous metrics of expression breadth and variability are provided for each ORF.

##### Supplemental Notes

**Supplemental Note N1: Theoretical Foundation of Hill Numbers and Perplexity.** We summarize the theoretical characterization of Hill numbers, and the special case of perplexity, developed in Leinster 2021. Hill numbers have a strong foundation in ecology for quantifying species diversity in ecological communities. It satisfies desirable mathematical properties including continuity, the replication principle, and modular-monotonicity. Perplexity is a Hill number of order 1 that satisfies desirable mathematical properties including continuity, the replication principle, and modular-monotonicity. These properties make perplexity particularly suitable for measuring isoform diversity.

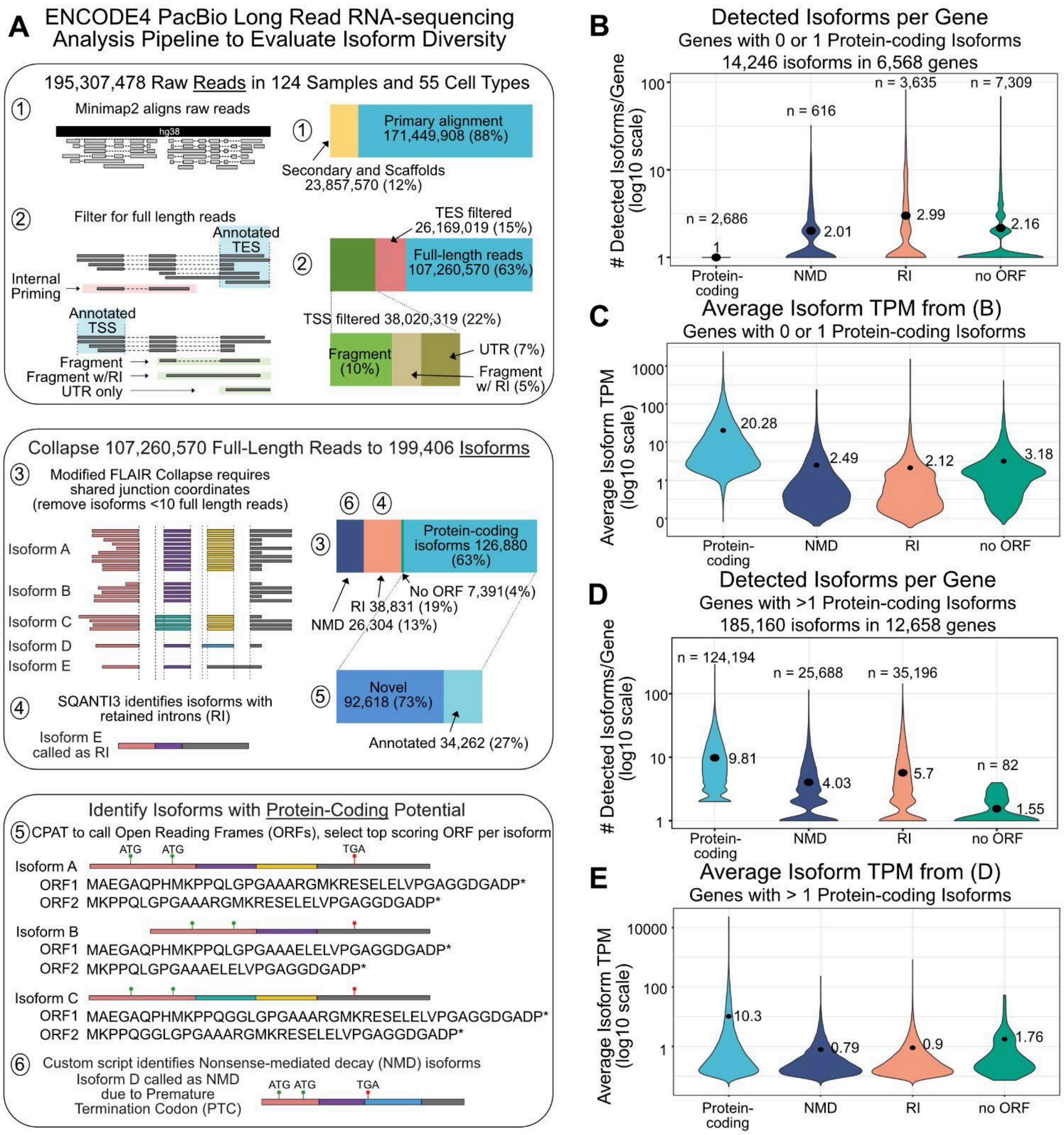


**Supplemental Figure 1.** (A) Schematic of our custom analysis pipeline to process ENCODE4 PacBio LRS data. Raw reads were aligned to the reference genome (hg38) using Minimap2 (Step 1), followed by filtering for full-length reads matching annotated transcription start and end sites (TSS and TES; Step 2). Isoforms were collapsed using a modification of the *FLAIR Collapse* script requiring shared splice junctions and a minimum of 10 full-length supporting reads across samples (Step 3). SQANTI3 was used to classify retained introns (RI), which were then removed (Step 4). CPAT was applied to identify open reading frames (ORFs; Step 5), and a custom script removed isoforms predicted to undergo nonsense-mediated decay (NMD; Step 6). Barplots to the right of each schematic show the number of reads or isoforms at each step of processing. The circled numbers 1-6 show step(s) to which each barplot corresponds. Percentages and counts are indicated for raw reads, full-length reads, and isoforms. (B) The number of detected isoforms per gene, for genes with 0-1 protein-coding isoforms, grouped by transcript biotype. NMD = nonsense-mediated decay; RI = retained intron. Numbers next to each violin are the average, and ‘n’ values above indicate the number of isoforms per biotype. (C) Averaged isoform TPMs across samples for genes in (B). Numbers next to the violins are the average TPM per transcript biotype. (D-E) Analogous plots to B-C but for genes with multiple (>1) protein-coding isoforms.

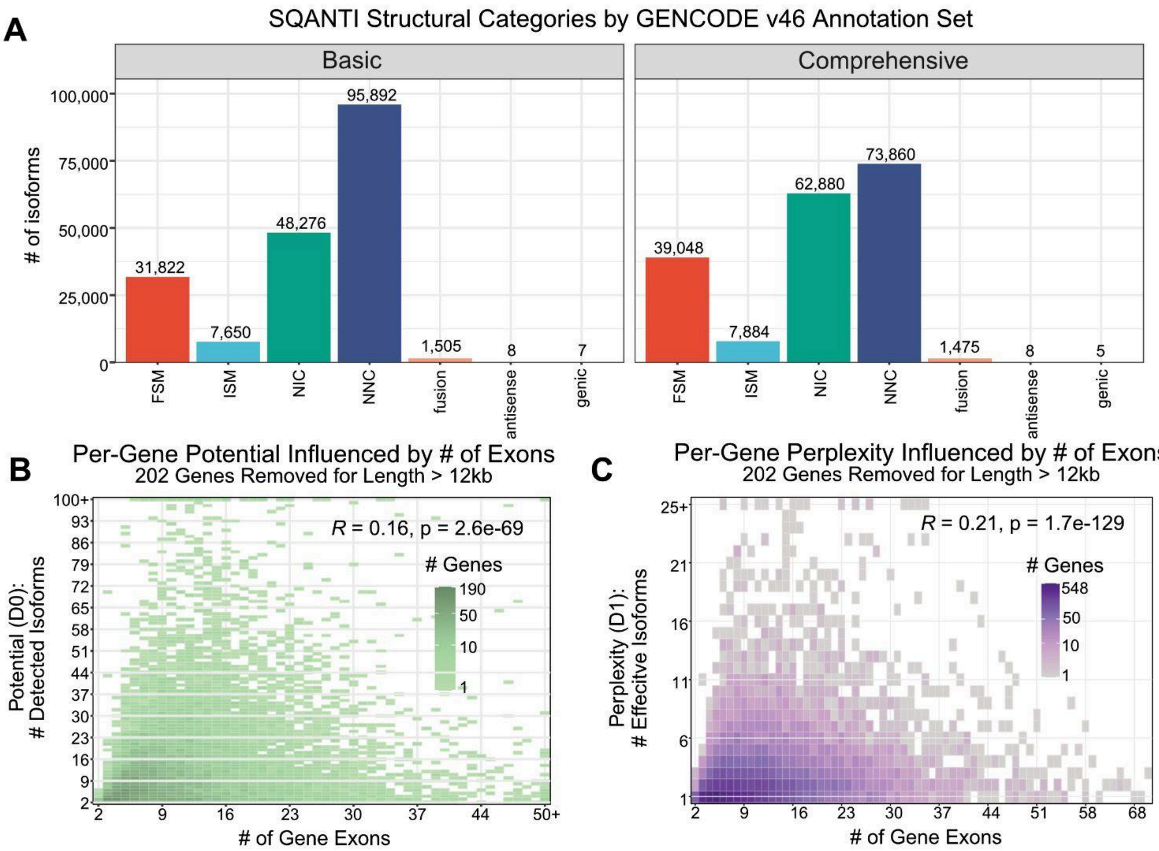


**Supplemental Figure 2.** (A) SQANTI3 structural categories for all 185,160 isoforms using GENCODE v46 basic (left) and comprehensive (right) annotations. FSM = full splice match, ISM = incomplete splice match, NIC = novel in catalog, NNC = novel not in catalog. The number of isoforms is shown above each bar. (B-C) Relationship between number of gene exons and per-gene potential (B) and per-gene perplexity (C). Genes with an annotated isoform longer than 12kb were excluded (n = 202) due to the known length bias in long-read sequencing. Color intensity corresponds to the number of genes at each point. Pearson's *R* and p-values are shown.

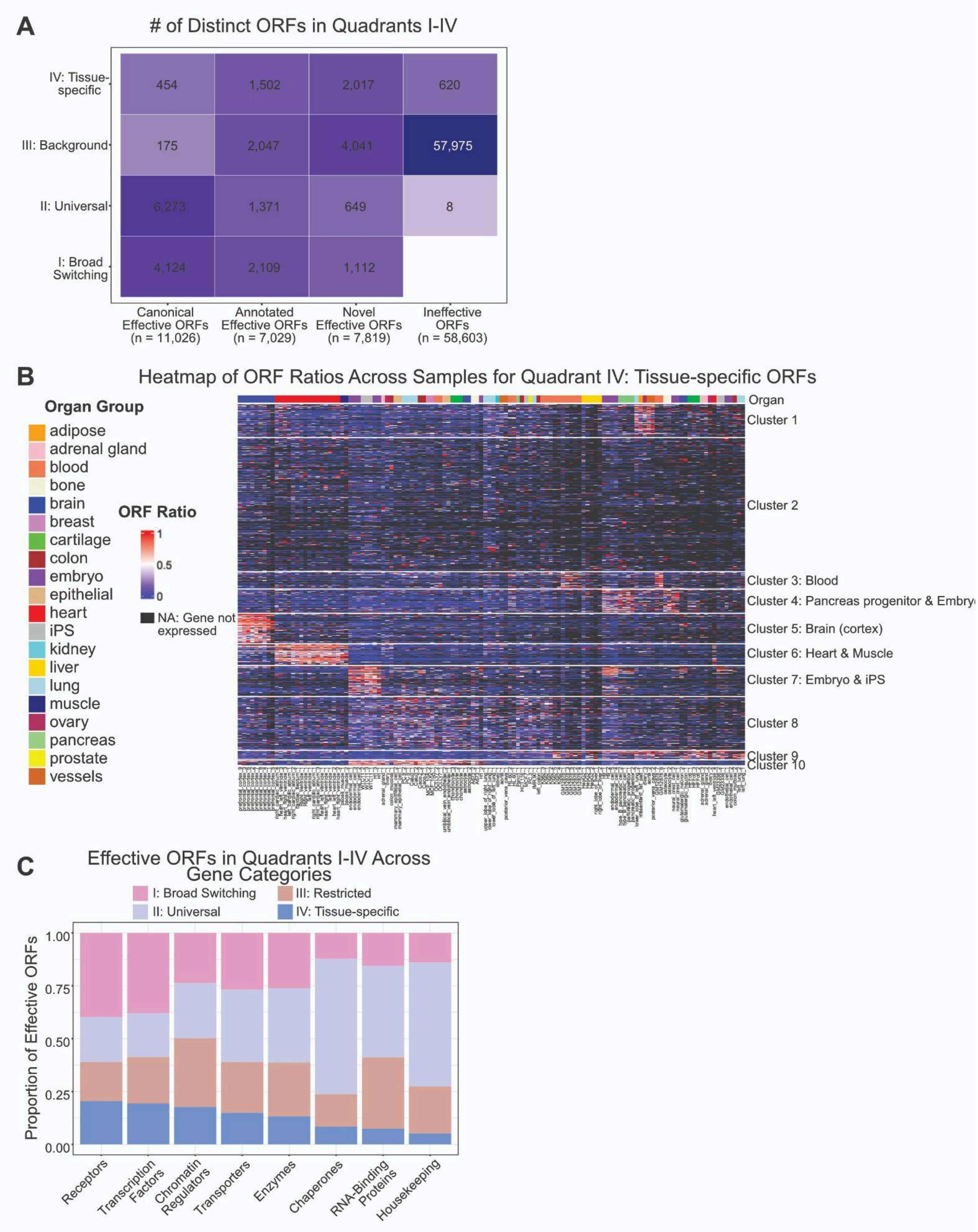
A
### of Distinct ORFs in Quadrants I-IV
IV: Tissue-specific
III: Background
II: Universal
I: Broad Switching
454
1,502
2,017
620
175
2,047
4,041
57,975
6,273
1,371
649
8
4,124
2,109
1,112
Canonical Effective ORFs (n = 11,026)
Annotated Effective ORFs (n = 7,029)
Novel Effective ORFs (n = 7,819)
Ineffective ORFs (n = 58,603)
B
Heatmap of ORF Ratios Across Samples for Quadrant IV: Tissue-specific ORFs
Organ Group
adipose
adrenal gland
blood
bone
brain
breast
cartilage
colon
embryo
epithelial
heart
iPS
kidney
liver
lung
muscle
ovary
pancreas
prostate
vessels
ORF Ratio
1
0.5
0
NA: Gene not expressed
Organ
Cluster 1
Cluster 2
Cluster 3: Blood
Cluster 4: Pancreas progenitor & Embry
Cluster 5: Brain (cortex)
Cluster 6: Heart & Muscle
Cluster 7: Embryo & iPS
Cluster 8
Cluster 9
Cluster 10
C
Effective ORFs in Quadrants I-IV Across Gene Categories
I: Broad Switching
II: Universal
III: Restricted
IV: Tissue-specific
Proportion of Effective ORFs
1.00
0.75
0.50
0.25
0.00
Receptors
Transcription Factors
Chromatin Regulators
Transporters
Enzymes
Chaperones
RNA-Binding Proteins
Housekeeping

**Supplemental Figure 3.** (A) The number of distinct ORFs across ORF types and quadrants I-IV. X-axis numbers are sums of columns. There are no ORFs that are both Broad Switching and Ineffective. (B) Tissue-specific ORFs are hierarchically clustered by Euclidean distance of their ORF usage ratios, resulting in ten groups. Organ types are marked at the top of the heatmap. For samples where the gene is not expressed, the ORF ratio is NA and colored black on the heatmap whereas a value of 0 indicates that the gene is expressed but the ORF is not. (C) The proportion of effective ORFs from each quadrant across the different gene categories–ordered on the x-axis by the proportion in type IV: Tissue-specific. For example, Receptors have the highest proportion of tissue-specific ORFs, while housekeeping has the lowest.
